# SOX2^+^ sustentacular cells are stem cells of the postnatal adrenal medulla

**DOI:** 10.1101/2023.10.28.564519

**Authors:** Alice Santambrogio, Yasmine Kemkem, Thea L. Willis, Ilona Berger, Maria Eleni Kastriti, Louis Faure, John P. Russell, Emily J. Lodge, Val Yianni, Rebecca J. Oakey, Barbara Altieri, Stefan R. Bornstein, Charlotte Steenblock, Igor Adameyko, Cynthia L. Andoniadou

## Abstract

Renewal of the catecholamine-secreting chromaffin cell population of the adrenal medulla is necessary for physiological homeostasis throughout life. Definitive evidence for the presence or absence of an adrenomedullary stem cell has been enigmatic. In this work, we demonstrate that a subset of sustentacular cells endowed with a support role, are in fact adrenomedullary stem cells. Through genetic tracing and comprehensive transcriptomic data of the mouse adrenal medulla, we show that cells expressing *Sox2/*SOX2 specialise as a unique postnatal population from embryonic Schwann Cell Precursors and are also present in the normal adult human adrenal medulla. Postnatal SOX2^+^ cells give rise to chromaffin cells of both the adrenaline and noradrenaline lineages *in vivo* and *in vitro.* We reveal that SOX2+ stem cells have a second, paracrine role in maintaining adrenal chromaffin cell homeostasis, where they promote proliferation through paracrine secretion of WNT6. This work identifies SOX2^+^ cells as a true stem cell for catecholamine-secreting chromaffin cells.

## Introduction

The adrenal medulla is responsible for the body’s reaction to acute stress and mediates the “fight or flight” response through the production and release of the catecholamines adrenaline, noradrenaline and low levels of dopamine, by specialised neuroendocrine chromaffin cells. Catecholamines target multiple organs to help increase oxygenation of muscles, blood pressure and heart output, blood sugar levels, attention and focus, and promote vasoconstriction as well as enhance memory performance ^1,2^. Diseases of the adrenal, such as congenital adrenal hyperplasia, dopamine beta-hydroxylase deficiency and tumours (pheochromocytomas and the related paragangliomas) lead to disruption in catecholamine regulation with life-threatening consequences ^3–5^.

The study of adrenal medulla homeostasis, the consequences of homeostatic perturbation, and prospects for regenerative approaches are lacking due to an incomplete characterisation of cell types in this organ. Previous *in silico* studies of the adrenal gland have delivered insights into adrenocortical cell transcriptome but failed to provide a characterisation of adrenomedullary cell subtypes. This is mostly due to cell isolation protocols being optimised for the adrenal cortex, resulting in very low viable cell numbers from the medulla ^6–8^. The existence of a putative adrenomedullary stem cell, capable of giving rise to new chromaffin cells *in vivo*, has previously been postulated but not identified ^9–11^. Instead, divisions in this slow-turnover organ have been attributed to chromaffin cells ^12,13^. The poor understanding of the adrenomedullary transcriptomic landscape has even hindered the discovery of specific cell markers within the heterogeneous chromaffin subtypes, such as the noradrenaline-producing chromaffin cells which, lacking a specific marker, are instead identified by the lack of expression of phenylethanolamine N-methyltransferase (PNMT), which catalyses the conversion of noradrenaline to adrenaline ^6–8^.

In this work, using single-cell RNA sequencing with cell isolation methods optimised for adrenomedullary cells, we identify a postnatal adrenomedullary stem cell population expressing the transcription factor SOX2. Through *in vivo* lineage tracing and *in ovo* assays, we demonstrate that these are a specialised long-lived population of stem cells originating and specialising from the embryonic precursors of the adrenal medulla, termed Schwann cell precursors (SCPs) ^14–16^. These adult SOX2+ stem cells contribute to the generation of new noradrenaline- and adrenaline-secreting chromaffin cells throughout life and promote organ proliferation through paracrine signalling. The identification of this novel adrenomedullary stem cell population holds promise for applications in regenerative medicine in neuroendocrine structures and constitutes an ideal target for oncogenic mutations known to cause pheochromocytomas and paragangliomas, which present some of the highest rates of gene heritability across all tumours.

## Results

### Transcriptomic analysis elucidates cell composition of the postnatal adrenal medulla and identifies a putative progenitor/stem cell population

To investigate the postnatal cell composition of the adrenal medulla, we performed droplet-based single-cell RNA sequencing on 10 mouse adrenals that were manually dissected to remove the majority of the cortex, at postnatal day (P) 15, during the rapid growth phase of the gland ^17^ (schematic Figure 1A) (*n* = 5 mice, mixed sex). Adrenomedullary cells were subset *in silico* using markers listed in Figure S1B, and all cortex, endothelial and immune cell types were excluded (Figure S1A, B). Following quality control (Figure S1C, D) unsupervised clustering of 2708 medullary cells revealed 8 distinct transcriptional signatures (Figure 1B,C). Cluster identity was assigned based on differential expression of known cell markers (Figure 1D).

**Figure 1.**
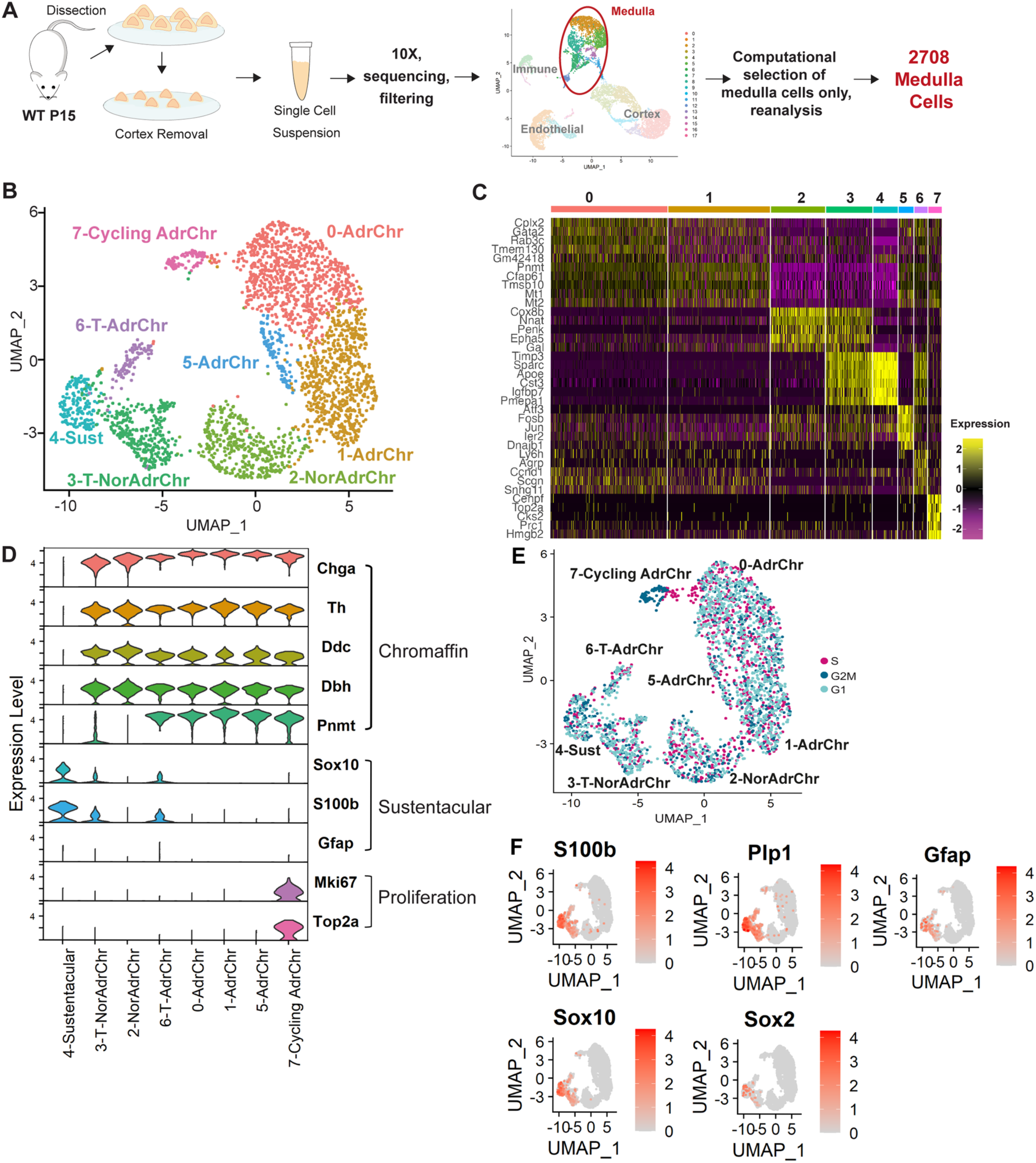
Single-cell RNA sequencing of the mouse adrenal medulla. A) Experimental workflow. B) UMAP of adrenomedullary cell types (2708 cells). C) Heatmap showing the transcriptional signatures of 8 clusters (top 5 differentially expressed genes). D) Violin plots indicating expression of different markers to identify each cluster. E) UMAP showing the distribution of cell cycle states over the dataset. Computationally identified percentages of medullary cells in different cell cycle phases: 44.61% (1208 cells) are in G1, 31.75% (859 cells) are in S, 23.67% (641 cells) in G2/M; F) Featureplots of known sustentacular cell and SCP markers *S100b*, *Plp1*, *Gfap*, *Sox10*, and of newly identified marker *Sox2*. Colour scale indicates expression level.

All chromaffin cells express tyrosine hydroxylase (TH), which converts tyrosine to L-DOPA, the precursor of dopamine. The action of dopamine-β-hydroxylase, encoded by *Dbh*, converts dopamine into noradrenaline, and subsequently, phenylethanolamine N-methyltransferase (PNMT) converts noradrenaline to adrenaline. Two types of chromaffin cells exist; a first type that expresses PNMT and secretes adrenaline, and a second type, present in the minority, not expressing PNMT and secreting noradrenaline. The differentiated chromaffin cells were further divided into 5 clusters: three consistent with adrenaline-producing signatures (Clusters 0 – 821 cells, 1 – 717 cells, 5 – 105) expressing *Chga*, *Th*, *Ddc*, *Dbh* and *Pnmt*, revealing heterogeneity amongst this population; one noradrenaline-producing (Cluster 2 – 382 cells) expressing *Chga*, *Th*, *Ddc* and *Dbh* but not *Pnmt*. In the literature, postnatal noradrenaline chromaffin cells were so far only recognised owing to the lack of *Pnmt* expression and were lacking identifying markers. Our study identifies three such unique markers expressed both within clusters 2 as well as the less-committed cluster 3, *Cox8b*, *Lix1* and *Penk* (Figure S1E). *Penk* encodes a preproprotein, whose products have previously been identified in chromaffin cell extracts ^18^. Immunofluorescence staining for PENK, demonstrates overlap with TH, marking all chromaffin cells, and mutually exclusive expression with PNMT, marking adrenaline-producing chromaffin cells (Figure S1F). We therefore consider PENK a reliable marker of noradrenaline chromaffin cells and further confirm its expression in human normal adrenal (Figure S1G). Consistent with previous literature ^12,13^, we additionally identify a fifth cluster of committed chromaffin cells, designated as cycling chromaffin cells (cluster 7 Cycling Chr – 92 cells) expressing chromaffin cell markers (including both *Pnmt* and *Penk*), as well as *Mki67* and *Top2a*. Cell cycle analysis confirmed that the majority of cluster 7 cells were in the G2M phase. Cells in G2M were also identified across all clusters, including sustentacular cells (Figure 1E).

We detected the presence of a cluster not expressing any chromaffin cell markers, indicative of possible progenitor/stem cells. This was designated as the sustentacular cell cluster, based on the known expression of markers of previously-described sustentacular cells, a signature partly shared by SCPs, the embryonic progenitors of the adrenal medulla. These included markers *Plp1*, *Lgi4*, *Fabp7*, *Sfrp1* and *Cdh19* ^14,19^ (Cluster 4 - 170 cells), (Figure 1F, S1H). The stem/progenitor markers *Sox10*, *S100b*, *Gfap* were expressed among this postnatal population, as well as *Sox2*, not previously reported but a marker of multiple progenitor/stem cells^20–23^ (Figure 1F). These genes all exhibited transcriptional heterogeneity amongst the sustentacular cell population and were additionally expressed, albeit at reduced expression levels, in two additional clusters. These were designated transitioning cell clusters (Clusters 3 T-NorAdrChr – 328 cells and 6 T-AdrChr – 93 cells), as they shared transcriptional signatures with both chromaffin and sustentacular cell markers and are likely committing progenitors of the two types of differentiated chromaffin cells (Figure 1D, S1I). This observation is supported by pseudotime inference, which predicts chromaffin cells arising from sustentacular cells via the transitioning clusters (Figure S1J). In summary, these data support the presence of a postnatal adrenomedullary progenitor/stem cell population within the previously termed sustentacular cells and indicate two branches of progenitors during commitment to either noradrenaline- or adrenaline-producing chromaffin cells.

### SOX2^+^ postnatal cells are a distinct SCP-derived subpopulation of sustentacular cells, present in the adrenal medulla throughout life

We next sought to determine if the expression of SOX2 marks a distinct subset of this sustentacular cell population. Immunohistochemistry using antibodies against SOX2 on sections of murine adrenals revealed that SOX2^+^ cells are present in the adrenal medulla during the early postnatal period and adulthood (Figure 2A). Quantification of SOX2^+^ cells as a proportion of the total cells in the medulla reveals an increase in SOX2^+^ cell proportion between P15 and P17 (4.95% to 7.04%), followed by a gradual decrease until P42 and maintenance of the proportion of SOX2^+^ cells until P365 (Figure 2B). We did not observe a difference in SOX2^+^ cell proportions between sexes (Figure S2A), of relevance since previous reports indicate a discrepancy in the volume of murine medulla based on sex ^17^. Immunostaining for SOX2 in human adrenals into advanced age (from patients aged 17, 29, 48, 55, 56, 71), confirms the presence of SOX2^+^ cells in the human adrenal medulla (Figure S2B). To validate overlap with sustentacular markers we used the *Sox2^eGFP/+^* mouse line where EGFP is expressed under the control of *Sox2* regulatory elements ^21^ and confirmed that all SOX2^+^ cells express EGFP (Figure S2C). Double-immunofluorescence staining confirms that SOX2^+^ cells express classical sustentacular cell markers SOX10, S100B and GFAP (Figure 2C). RNAscope mRNA *in situ* hybridisation confirms transcripts of both *Sox2* and *S100b* or *Gfap* in the same cells and additionally reveals overlap of *Sox10* and *Plp1* with *Sox2*, affirming the shared signature with SCPs (Figure 2D). Analysis of a published developmental SCP dataset (Kastriti et al. 2022) reveals that *Sox2* is expressed among ‘multipotent hub’ cells, and that *Sox10* expression precedes *Sox2* expression in the chromaffin cell commitment trajectory (Figure 2E). The *Sox2* regulon is active in the uncommitted state of the chromaffin commitment trajectory (Figure S2D). The top 10 markers correlated with *Sox2* expression across all cells include genes highly expressed in postnatal sustentacular cells (*Fabp7*, *Sparc*, *Zfp36l1*, *Serpine2 Sox10*), and the Hippo pathway regulator *Wwtr1* (Figure S2E and F). Anti-correlated genes (analysis only among *Sox2*-expressing cells) include chromaffin cell markers *Chga*, *Chgb* and *Th* (Figure S2G), supporting the notion that *Sox2* expression needs to be downregulated for acquisition of a chromaffin cell state. To determine if these SOX2^+^ sustentacular cells are indeed derived from SCPs in the embryo, we carried out lineage tracing of embryonic SCPs. Using *Wnt1^Cre/+^;R26^mTmG/+^* we labelled the neural crest from its specification and using *Sox10^iCreERT^*^2^*^/+^*;*R26^mTmG/+^* we induced SCPs at 11.5dpc, at a time when they begin to migrate towards the dorsal aorta, subsequently giving rise to the adrenal medulla. Immunofluorescence staining using GFP and SOX2 antibodies, reveals that descendants of *Wnt1* and *Sox10* expressing cells (GFP+) include the entire SOX2^+^ population at P15 (Figure 2F, G). In order to determine if postnatal SOX2^+^ cells are a distinct specialised cell type from SCPs, we compared the signatures of SCPs and postnatal SOX2^+^ cells. Isolation of a flow-purified population enriched for SOX2^+^ cells using the *Sox2^eGFP/+^* mouse line, allowed single-cell RNA sequencing of 1563 cells, 493 of which are expressing high levels of *Sox2* (Figure S3A). Comparison of the transcriptomic signature of *Sox2*^High^-expressing subset to SCPs identifies that postnatal SOX2^+^ cells have a distinct signature, supporting that these have become a specialised population of progenitor/stem cells (Figure S3B, C). STRING analysis of the top differentially expressed genes reveal that SOX2^+^ postnatal cells differentially express a hub of extracellular matrix-related genes including *Col4a1*, *Col4a2*, *Col28a1*, *Col26a1* and a hub signalling/transcription factors including *Fos*, *Fosb*, *Egr1*, *Ier2*, *Klf2*, *Ctgf* (Figure S3D). In summary, *Sox2*-expressing cells of the postnatal adrenal medulla are derived from SCPs and are a distinct uncommitted postnatal population.

**Figure 2.**
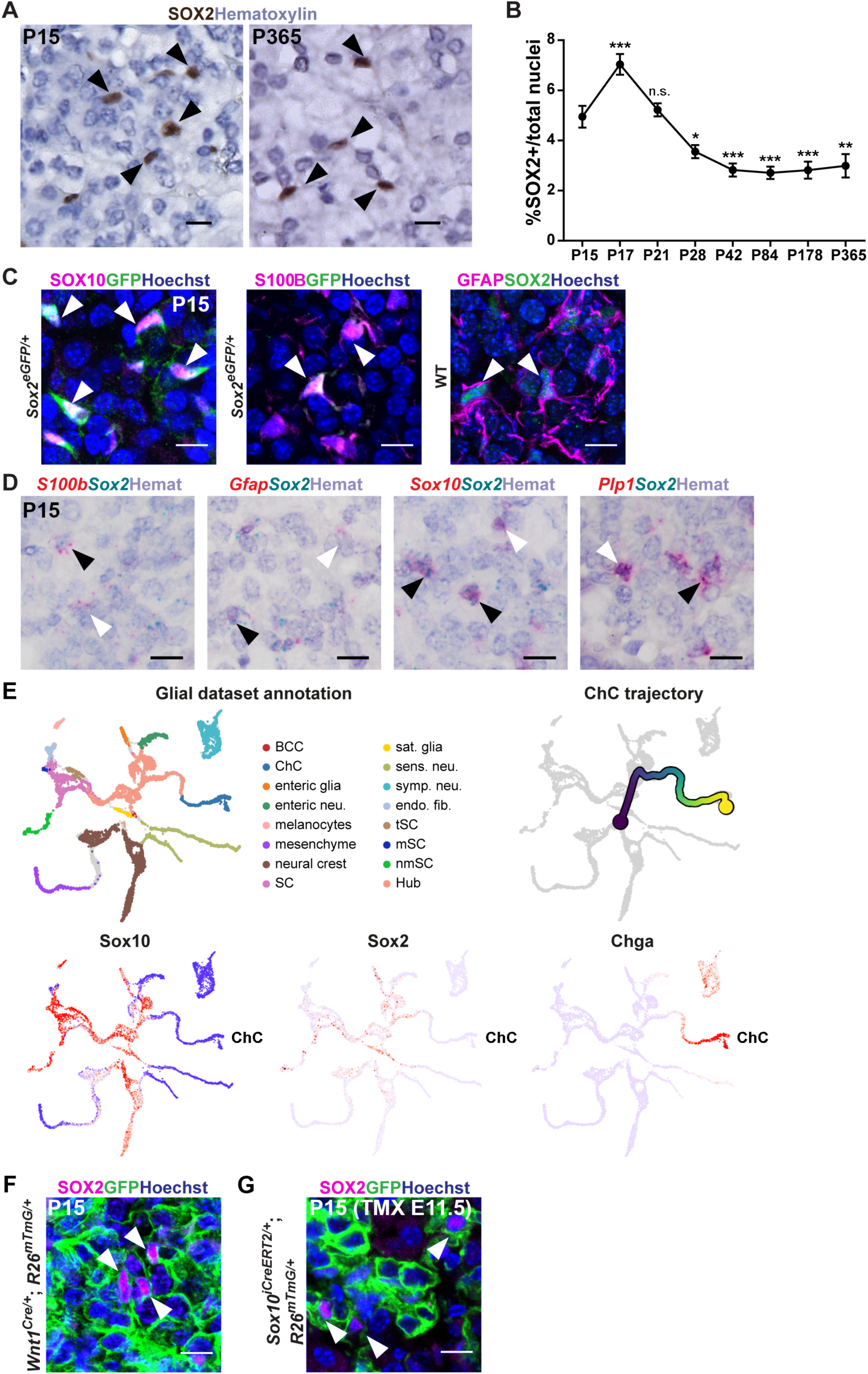
SOX2^+^ cells are present in the adrenal medulla and are derived from Schwann Cell Precursors. A) Immunohistochemistry with antibodies against SOX2 (brown) at P15 and 365 in wild type adrenal medullae. Nuclei counterstained with Hematoxylin, scale bar 20μm. B) Quantification of SOX2^+^ cells over the total nuclei of adrenal medulla. n = 6 animals for each timepoint, plotted mean and SEM. One-way ANOVA multiple comparisons test: P15 vs. P17 (*P*-value 0.0009); P15 vs. P21 (*P*-value 0.9925); P15 vs. P28 (*P*-value 0.0386); P15 vs. P42 (*P*-value 0.0007); P15 vs. P84 (*P*-value 0.0003); P15 vs. P178 (*P*-value 0.0006); P15 vs. P365 (*P*-value 0.0013). C) Immunofluorescence staining of P15 *Sox2^eGFP/+^* adrenal medulla using antibodies against SOX10 (magenta) or S100β (magenta) and GFP (green), shows double-positive cells in both (arrowheads). Immunofluorescence staining of a P15 wild type (WT) sample using antibodies against GFAP (magenta) and SOX2 (green) shows double-positive cells (arrowheads). Nuclei counterstained with Hoechst, scale bars 10μm. D) RNAscope mRNA *in situ* hybridisation on wild type P15 samples shows double-positive cells for *S100b* (red) and *Sox2* (blue), *Gfap* (red) and *Sox2* (blue), *Sox10* (red) and *Sox2* (blue), *Plp1* (red) and *Sox2* (blue) respectively - (black arrowheads) or single positive (white arrowheads); all nuclei counterstained with Hematoxylin, scale bar 10μm. E) UMAP of the neural crest and SCP lineages between 9.5dpc - 12.5dpc from ^29^. Trajectory of chromaffin cell (ChC) transitioning from the Hub cells. Featureplots showing expression of *Sox10*, *Sox2*, *Chga*. *Sox10* expression precedes that of *Sox2* in this trajectory, which is maintained until expression of *Chga* marking chromaffin cells. F) Immunofluorescence on P15 mouse adrenals from *Wnt1^Cre/+^;R26^mTmG/+^*genotypes. Immunostaining with antibodies against SOX2 (magenta) and GFP (green) shows double-positive cells (arrowheads). Nuclei are counterstained with Hoechst, scale bars 10μm. G) Immunofluorescence on P15 mouse adrenals from a *Sox10^CreERT^*^2^*^/+^;R26^mTmG/+^* line induced with tamoxifen (TMX) at 11.5dpc – immunostaining with antibodies against SOX2 (magenta) and GFP (green) shows double-positive cells (arrowheads). Nuclei counterstained with Hoechst, scale bars 10μm.

### Isolated Sox2-expressing stem cells self-renew and give rise to new chromaffin cells ex vivo

To determine if *Sox2*-expressing cells have the potential for self-renewal, we cultured dissociated adrenomedullary cells in stem-cell promoting media under adherent conditions. Flow sorting of EGFP^+^ and EGFP^-^ cells from *Sox2^eGFP/+^* mice at P15 and plating in clonogenic conditions, showed that EGFP^+^ (SOX2^+^) cells only, can give rise to colonies, which can be passaged (Figure 3A, B, S4A-C). These colonies contained an expanded population of SOX2^+^ cells as revealed by immunofluorescence staining and by flow cytometry for EGFP expression (Figure 3C, D). These data combined, render *Sox2*-expressing cells as a putative postnatal stem cell population. To establish if SOX2^+^ cells alone are sufficient to give rise to chromaffin cells, we took advantage of a well-established *in vivo* xenograft culture technique, chorioallantoic membrane (CAM) culture, in chicken embryos ^24^. This allows culture of three-dimensional vascularised tissues in an *in vivo* environment, enabling long-term maintenance. We first used our newly-established *in vitro* culture system to isolate and expand postnatal SOX2^+^ stem cells over 8 days, at which point they are mostly uncommitted, as revealed by EGFP detection by flow sorting (Figure 3D). Purified SOX2^+^ cell suspensions (800,000 cells, purified and expanded from 4-6 animals) were grafted onto the embryonic CAM (Figure 3E). Collection of the xenografts 10 days later, revealed that SOX2^+^ cells can give rise to compact three-dimensional tissues (*n=4* out of 10 CAM assays, Figure 3F). Endogenous expression of EGFP was detectable in the grafts at collection, suggesting the presence of SOX2^+^cells (Figure 3F). Immunofluorescence staining using antibodies against chromaffin cell markers TH and PNMT (adrenaline-expressing chromaffin cells), confirms that grafts contain differentiated chromaffin cells (*n=3* grafts, Figure 3G).

**Figure 3.**
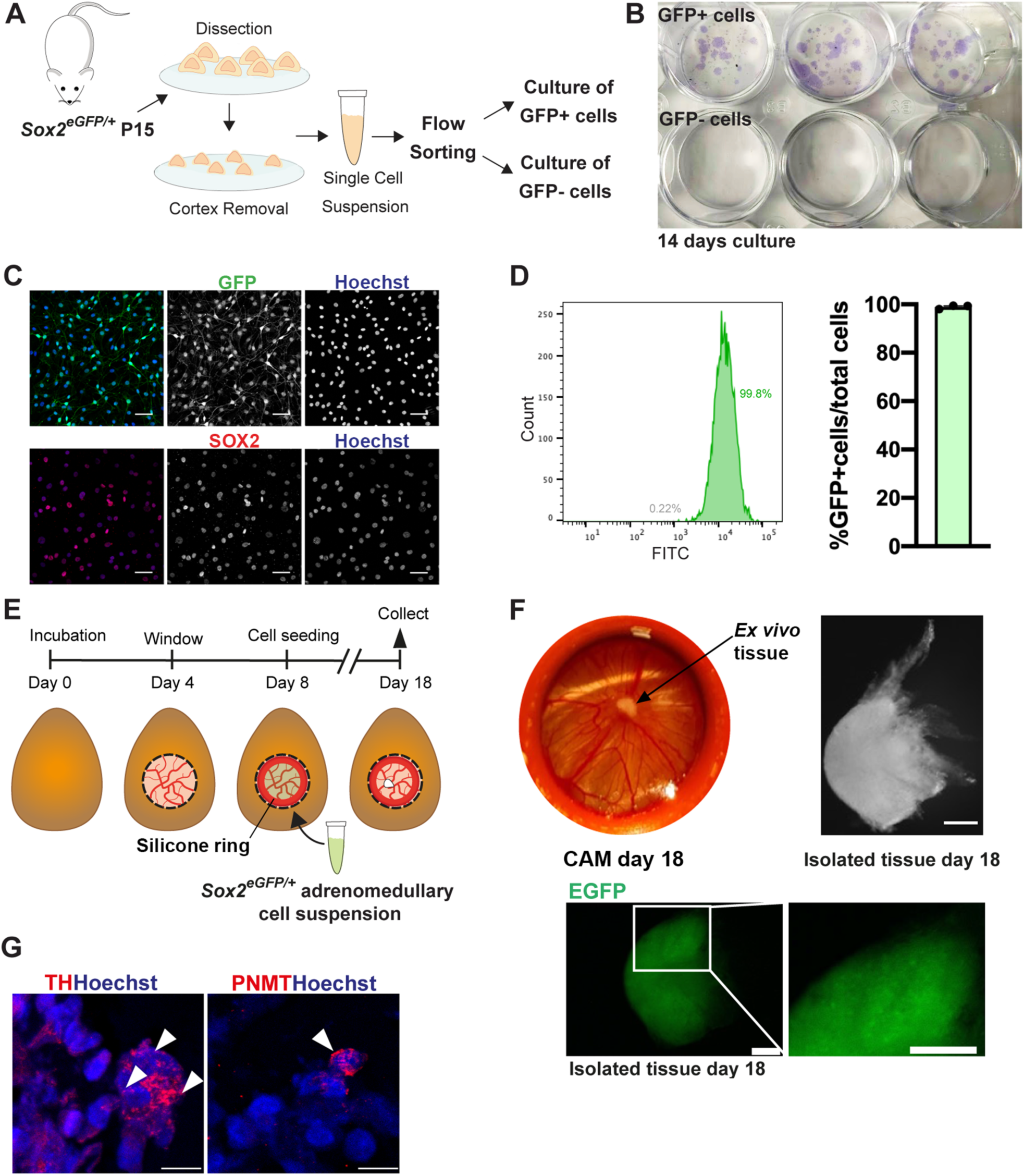
Adrenomedullary SOX2^+^ cells have stem cell properties *in vitro* and *in ovo*. A) Experimental workflow. B) Crystal violet staining of fixed cell colonies following 14 day culture of GFP^+^ (SOX2^+^) and GFP^-^ (SOX2^-^) *Sox2^eGFP/+^* cells under clonogenic conditions. C) Immunofluorescence staining of GFP^+^ primary cells from *Sox2^eGFP/+^* medulla cultured for 14 days: GFP (green), SOX2 (red), nuclei stained with Hoechst, scale bar 50μm. D) Quantification of GFP^+^ cells via flow cytometry after 14 days of culture, bar graph *n*=3 independent biological replicates. E) Experimental design of chick chorioallantoic membrane (CAM) assays for *ex vivo* 3D xenograft culture. F) Representative images of resulting xenograft before removal from CAM (left) and after isolation (right) following 18 days of incubation. Representative images of wholemount native EGFP expression in *Sox2^eGFP/+^*-derived xenograft (bottom). Scale bars 200μm G) Immunofluorescence staining using antibodies against TH (red) or PNMT (red) on xenografts at day 18. Nuclei counterstained with Hoechst. Scale bars 10μm.

### Postnatal SOX2^+^ cells of the adrenal medulla are stem cells in vivo

To establish if SOX2^+^ cells function as stem cells during homeostasis *in vivo*, we labelled and lineage-traced *Sox2*-expressing cells in the postnatal adrenal. Tamoxifen induction of *Sox2^CreERT^*^2^*^/+^; R26^mTmG/+^* animals was carried out by single injection at P14 and adrenals collected after 72h (P17), 7 days (P21), 14 days (P28), 28 days (P42), 70 days (P84) and 178 days (P192) (Figure 4A). No-Cre controls or injection with corn oil instead of tamoxifen, did not result in GFP expression (Figure S4D). Lineage tracing over 178 days reveals an expansion of GFP^+^ clones in the adrenal medulla (Figure 4B). Quantification of GFP^+^ as a proportion of the total nuclei demonstrates an increase over time: initial labelling of 3.4% GFP^+^ cells at 72h post-induction to 10.67% at 178 days post-induction (Figure 4C). Double-immunofluorescence staining with antibodies against GFP and general chromaffin marker TH, adrenaline-producing chromaffin marker PNMT and the newly identified noradrenaline-producing chromaffin marker PENK, confirms GFP^+^ cells double-stained with either marker, confirming the derivation of both adrenaline- and noradrenaline-producing chromaffin cells from SOX2^+^ sustentacular progenitors (Figure 4D). Lineage tracing in adult mice induced at P189 (27 weeks n=4) and analysed 28 days later (P217), reveals the *in vivo* potential of *Sox2*-expressing cells to generate chromaffin cells is retained in later life (*n=3*, Figures 4E). Taken together, our data therefore demonstrate that SOX2^+^ adrenomedullary cells are *bona fide* stem cells.

**Figure 4.**
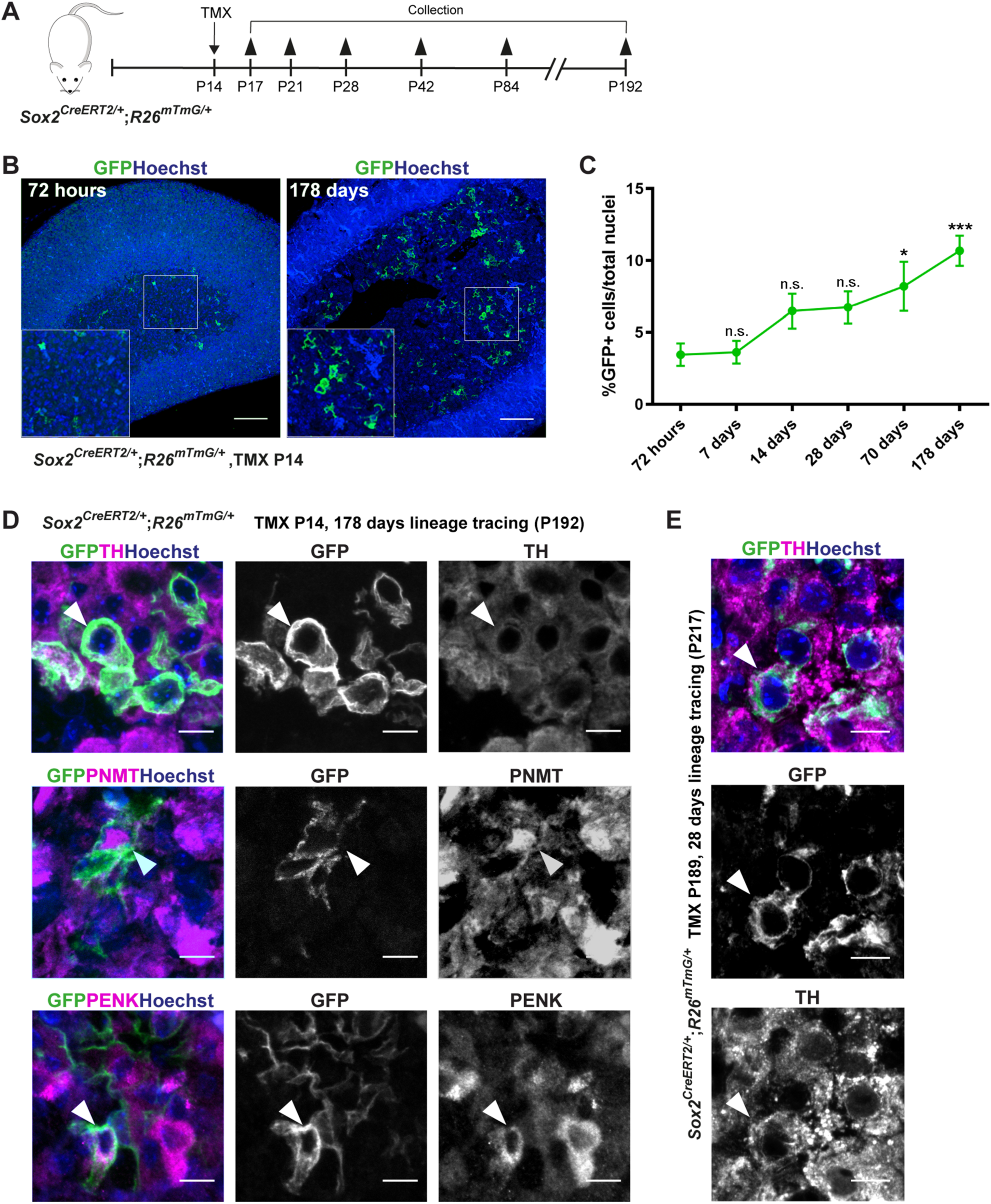
Adrenomedullary SOX2^+^ cells are stem cells *in vivo*. A) Experimental design indicating tamoxifen (TMX) induction at P14 and timepoints of collection and analysis. B) Immunofluorescence using antibodies against GFP, on *Sox2^CreERT^*^2^*^/+^;R26^mTmG/+^* adrenals induced with tamoxifen at P14 and collected after 72 hours or 178 days. GFP in green, nuclei counterstained with Hoechst. Inserts magnified boxed regions. Scale bar 100μm. C) Quantification of GFP+ cells/total nuclei of adrenal medulla at different timepoints. *N*=6 animals for each timepoint, plotted mean and SEM. One-way ANOVA multiple comparisons test: 72hrs vs. 7 days (*P*-value 0.9999); 72hrs vs. 14 days (*P*-value 0.2522); 72hrs vs. 28 days (*P*-value 0.1912); 72hrs vs. 70 days (*P*-value 0.0286); 72hrs vs. 178 days (*P*-value 0.0006). D) Double-immunofluorescence on *Sox2^CreERT^*^2^*^/+^;R26^mTmG/+^* adrenals induced at P14 and collected after 178 days, using antibodies against GFP (green) and specific cell markers (magenta) TH (all chromaffin cells), PNMT (adrenaline chromaffin cells) or PENK (noradrenaline chromaffin cells). Note the presence of double-labelled cells (arrowheads). Nuclei counterstained with Hoechst (blue), scale bar 10μm. E) Double-immunofluorescence on *Sox2^CreERT^*^2^*^/+^;R26^mTmG/+^* mice induced at P189 (6 months) and collected after 28 days, using antibodies against GFP (green) and TH (magenta). Note the presence of double-labelled cells (arrowheads). Nuclei counterstained with Hoechst (blue), scale bar 10μm.

### SOX2^+^ adrenomedullary stem cells produce paracrine WNT signals that promote expansion of surrounding endocrine cells

We previously reported that SOX2+ stem cells of a different endocrine organ, the anterior pituitary gland, are instrumental to promote postnatal organ proliferation in a paracrine manner, through the secretion of WNT ligands ^25^. To determine if SOX2+ stem cells of the adrenal medulla share this non-classical stem cell contribution to organ turnover, we mined our single-cell RNA s9equencing dataset of the mouse adrenal medulla to first explore the cell types that upregulate the WNT pathway. We found that WNT pathway targets *Lef1* and *Axin2* are both expressed in chromaffin cells, with a bias for the noradrenaline (*Lef1*) and adrenaline (*Axin2*) lineages. *Lgr5*, a WNT pathway potentiator and target is strongly expressed in all committed chromaffin cell clusters. All three targets are expressing in the cycling chromaffin cell cluster (cluster 7) (Figure 5A). Upregulation of canonical WNT signalling in chromaffin cells was confirmed through immunofluorescence against TH on the TCF/Lef:H2B-GFP reporter line, showing activation of GFP (WNT-responding cells) among TH+ chromaffin cells (Figure 5B). Expression of all three WNT targets was absent from the sustentacular/stem cell cluster. To determine the source of WNT ligands we queried expression of all mouse *Wnt* genes. *Wnt1*, *2*, *2b*, *3a*, *5b*, *7a*, *7b*, *8a*, *8b*, *9a*, *9b*, *10a*, *10b*, and *11* were not expressed in any adrenomedullary cell population. Low expression of *Wnt3*, *4*, *5a* and *16* was detected in sporadic cells across different clusters (Figure S5A). Expression of *Wnt6* was strong in the sustentacular/stem cell cluster, and detectable but weak in the two transitioning noradrenaline and adrenaline clusters (Figure 5C). Expression of *Wnt* genes within the isolated SOX2-EGFP^+^ cells confirmed robust expression of *Wnt6* as the sole *Wnt* gene (Figure 5C). Double mRNA in situ hybridisation using probes against *Wnt6* and *Sox2*, confirms specific expression of *Wnt6* in this stem cell population (Figure 5D, *n*=3). WLS (GPR177) is a glycoprotein receptor necessary for WNT secretion. *Wls* expression was detected across all populations of the adrenal medulla including the SOX2^+^ cells (Figure 5E). We specifically deleted *Wls* in SOX2^+^ cells (*Sox2^CreERT^*^2^*^/+;^Wls^fl/fl^*) in a tamoxifen-inducible manner. Mice were induced at P21 and adrenals collected at P26. Immunofluorescence staining using antibodies against Ki-67 to mark cycling cells, revealed a reduction in overall proliferation in adrenal medullae deficient in *Wls* expression (lack of WNT secretion) from SOX2^+^ cells (Figure 5G, *n*=6-8 per genotype). These results confirm that SOX2^+^ cells promote proliferation in the adrenal medulla, through paracrine secretion of WNT6.

**Figure 5.**
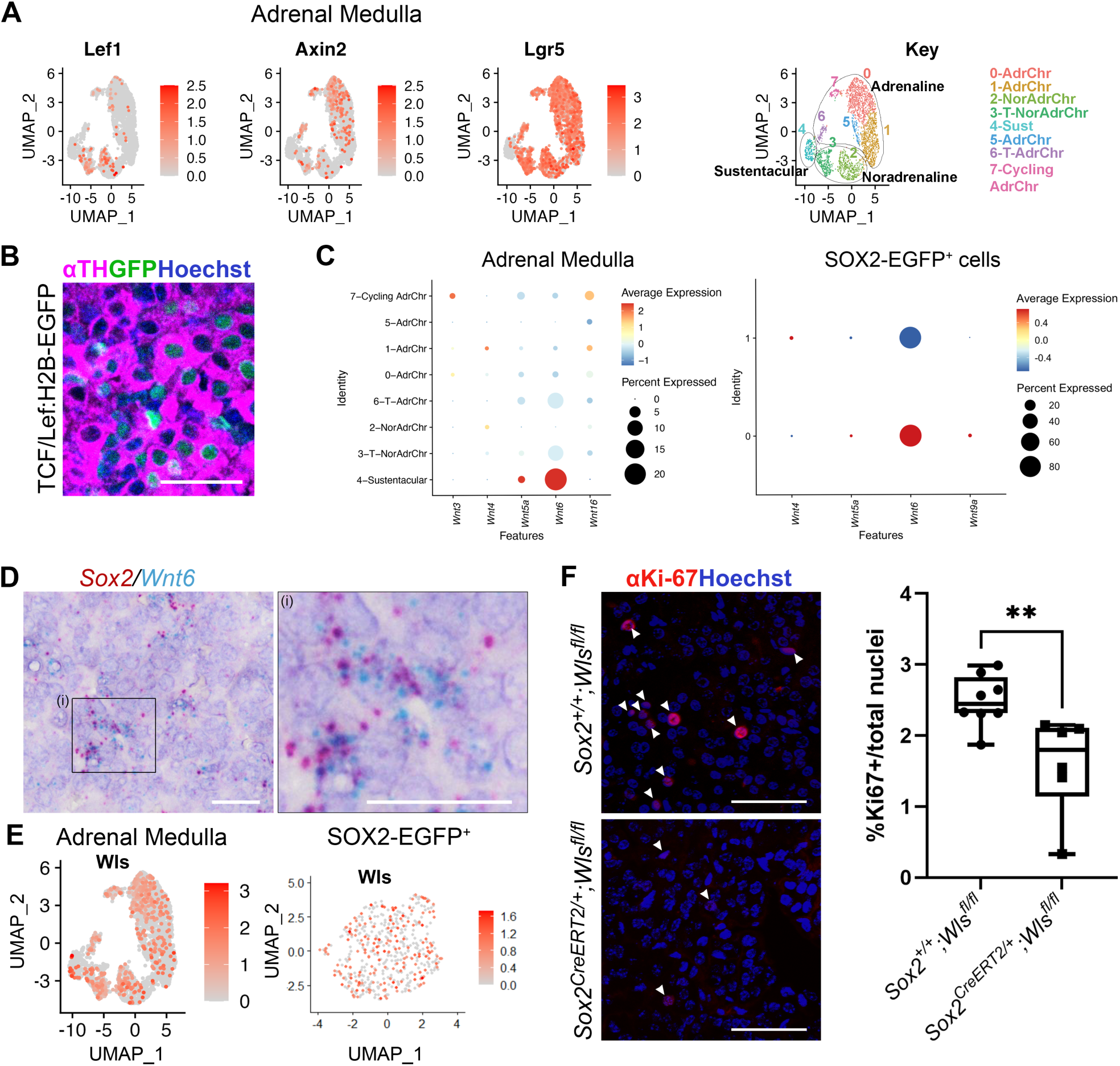
SOX2+ adrenomedullary stem cells promote proliferation of chromaffin cells through secretion of paracrine WNT signals. A) Featureplots for expression of WNT targets *Lef1*, *Axin2* and *Lgr5* in the mouse postnatal adrenal medulla dataset, Key of clusters, grouping by lineage. B) Immunofluorescence using antibodies against TH (chromaffin cells) and GFP (cells that have responded to WNT) on mouse TCF/Lef:H2B-EGFP adrenal medulla at P21. Nuclei are counterstained with Hoechst, scale bars 50 μm. C) Dot plots of all *Wnt* genes expressed in the mouse adrenal medulla dataset (left) and in the isolated SOX2-EGFP^+^ cell dataset (right), subdivided by cell clusters. D) RNAscope mRNA in situ hybridisation using probes against *Sox2* (red) and *Wnt6* (blue), showing co-expression. Image in right is magnified region (i). Scale bars 20μm. E) Featureplots for *Wls* in the mouse adrenal medulla dataset (left) and in the isolated SOX2-EGFP^+^ cell dataset. F) Representative immunofluorescence using antibodies against Ki-67 marking cycling cells in *Sox2^+/+^;Wls^fl/fl^*(control, top) and *Sox2^CreERT^*^2^*^/+^;Wls^fl/fl^*(mutant, bottom) samples following tamoxifen induction at P21 and analysis at P26 (*n*=8 controls, 6 mutants). Nuclei counterstained with Hoechst, scale bars 50μm. Graph showing percentage of Ki-67 positive cells across replicates, revealing a statistically significant reduction in cycling cells in the mutant. Unpaired *t*-test, *P*-value = 0.0083.

## Discussion

Here, we reveal the existence of postnatal adrenomedullary stem cells, which give rise to new chromaffin cells of both the adrenaline and the noradrenaline lineages throughout life, as well as contribute to the normal turnover of chromaffin cells through paracrine signalling. Employing *in vivo* studies in mouse, we confirm that this specialised *Sox2*-expressing somatic stem cell population derives from *Sox10*-expressing embryonic Schwann cell precursors of the neural crest and becomes a stem cell population distinct from SCPs. Comprehensive single-cell transcriptome analyses of the murine adrenal medulla were lacking from the literature, since methodologies to dissociate adrenal tissue favour the cortex, with low medullary cell survival. These reveal the molecular features of adrenomedullary stem cells, and clearly identify them as the cells of origin of noradrenaline- and adrenaline-expressing chromaffin subtypes, with distinct transitioning progenitors. Genetic lineage tracing of *Sox2*-expressing cells, confirms the generation of new chromaffin cells of both types throughout postnatal life. The transcriptomic datasets presented can be exploited further by the community, and as proof-of-concept, we present their use for identification of noradrenaline-specific markers, where previously noradrenaline-secreting chromaffin cells were identified only through their lack of marker expression. Validation of PENK as a marker of noradrenaline cells in mouse and human is demonstrated, which can be useful in human pathology, the study of noradrenaline cells and expansion of the genetic toolkit of mouse models in adrenal research (e.g. *Penk-Cre* mice) ^26^.

Previously, it was unknown if an adrenomedullary stem cell population exists or if new chromaffin cells are only generated from self-duplication ^27^. In this study, we not only demonstrate the potential of SOX2^+^ cells *in vivo*, but show that they can be cultured and expanded *in vitro* and generate tissue containing neuroendocrine cells when explanted, here illustrated using an *in ovo* system. The culture systems we established can be further exploited for stem cell-based regenerative approaches in relation to disorders implicating the adrenal medulla e.g. adrenal hypoplasia or dopamine β-hydroxylase deficiency.

In addition to the classical stem cell paradigm, our *in vivo* results reveal that SOX2^+^ cells can promote turnover in a second way, through the secretion of paracrine ligands, and we identify WNT6 as a key ligand in this process. The paradigm of stem cells acting as signalling hubs to regulate proliferation of their neighbouring descendants was previously demonstrated in pituitary gland stem cells ^25^, and further shown to underlie tumour formation ^20^. It supports the possibility that adrenomedullary stem cells may contribute cell autonomously and cell non-autonomously to adrenal tumour pathogenesis. Since an established stem cell population had not been identified, the current dogma dictates the cell-of-origin of pheochromocytoma and the related paraganglioma tumours as being specialised neuroendocrine cells ^28^. It can be postulated that this newly characterised stem cells may be involved in the initiation or progression of tumours, and our findings can support study of these processes and the generation of disease models.

## Acknowledgements

We thank Professor Martin Fassnacht for support with human adrenal tissue, Professor Karen Liu and Dr William Barrell for support with CAM assays, the King’s College London Biological Services facilities, the Advanced Cytometry Platform (Flow Core), Guy’s and St. Thomas’ NHS Trust and the Genomics Research Platform, R&D Department, Guy’s and St. Thomas’ NHS Trust. We thank Prof. Françoise Helmbacher and Dr Marika Charalambous for critical reading of the manuscript and insightful comments.

## Author Contributions

Conceptualisation, A.S., C.L.A.; Methodology, A.S., Y.K., C.L.A.; Software and Formal Analysis, A.S., T.L.W., V.Y., L.F., I.A.; Investigation, A.S., Y.K., T.L.W., I.B., M.E.K., L.F., J.P.R., E.J.L.; Writing – Original Draft, A.S., Y.K., C.L.A.; Writing – Review & Editing, A.S., Y.K., T.L.W., I.B., M.E.K., L.F., J.P.R., E.J.L, V.Y., R.J.O., P.A., S.R.B., C.S., I.A., C.L.A. Funding Acquisition, C.L.A., R.J.O., S.R.B., C.S.; Resources, B.A., I.A.; Supervision, R.J.O., S.R.B., C.S., I.A., C.L.A.

## Ethics

Studies using human adrenals were carried out under King’s College London ethical approval with KCL Ethics Reference LRS-19/20-20118, as part of the adrenal tumour registry project of the European Network for Adrenal Tumours ENS@T (European Network for the Study of Adrenal Tumours). All animal studies were performed under compliance of the Animals (Scientific Procedures) Act 1986, Home Office Licences P5F0A1579 (mouse) and P8D5E2773 (chicken), as well as KCL Biological Safety approval for project ‘Function and Regulation of Adrenal Stem Cells in Mammals’.

## STAR Methods – Santambrogio *et al*

### Animals

Procedures were carried out in compliance with the Animals (Scientific Procedures) Act 1986, Home Office licence and King’s College London ethical review approval. All mouse colonies were maintained under 12:12 hours light/dark cycle and fed *ad libitum*. All mouse lines used were previously described: *Sox2^eGFP/+^* ^21^, *Sox2^CreERT^*^2^*^/+^* ^20^, *Wnt1^Cre/+^* ^30^, *Sox10^iCreERT^*^2^*^/+^* ^31^*, R26^mTmG/+^* ^32^. All mice were bred and maintained on mixed backgrounds and consistently backcrossed on CD1. For Cre recombination, Tamoxifen (Sigma, T5648) was injected intraperitoneally with a single dose of 0.15mg/g body weight in postnatal mice. Pregnant females were injected by a single intraperitoneal injection of tamoxifen, capped at 1.5mg and one dose of Progesterone (Sigma P0130) at 0.75mg.

### Human samples

Normal adrenal human tissue samples were obtained from the University Hospital Würzburg (Germany). Normal adrenal glands removed as part of tumour nephrectomy and proven to be histologically normal, showing no neoplastic tissue.

### Fluorescent Activated Cell Sorting

Adrenal glands from *Sox2^eGFP/+^* mice were dissected and tissue was dissociated as described for primary cell culture. At the last step, cells were resuspended in FACS buffer (2.5% HEPES solution 1M (Sigma), 1% FBS in PBS), passed through a 40μm cell strainer (Corning, CLS431750) and stained with DAPI 0.05μg/ml (Biolegend, 422801), before being flow sorted by a FACSAria Cell Sorter (BD Biosciences). Adrenal medullae from wild type littermates were used as a negative control.

### 10x Single-cell RNA sequencing and computational analysis - postnatal adrenal medulla

10 adrenals from 5 mixed sex P15 mice were dissected on ice, surrounding fat and excess adrenal cortex were removed manually. Medullae were placed in an enzymatic digestion mix containing 50μg/ml DNAse I (Sigma, D5025), 10mg/ml Collagenase II (Worthington, LS004177), 2.5μg/ml Fungizone (Gibco, 15290026), 0.1X Trypsin-EDTA (Sigma, 59418C) in 1X Hank’s Balanced Salt Solution (HBSS) (Gibco, 14025050) and incubated at 37°C for 15 minutes. Enzymes were inactivated by addition of 10 times volume of serum-containing Base Media: DMEM/F-12 (Gibco, 31330-038) + 5% FBS (Merk, F0804) + 50U/ml Penicillin-Streptomycin (Gibco, 15070063). The cell suspension was centrifuged at 281g for 5 minutes at room temperature, washed twice in PBS and pellets were resuspended in a solution of HBSS 2.5% FBS. An aliquot of 10,000 viable cells were used for the experiment. Library preparation and sequencing were performed by the BRC Genomics Core at KCL. Library preparation was done using the Chromium 10X Single-cell 3’ Reagent Kit v3.1 (10x Genomics, PN-1000121) and a Chromium Controller (10x Genomics) following the manufacturer’s protocol. Once obtained, barcoded transcripts from single cells were sequenced with an Illumina HiSeq 2500.

Pre-processing of the sequencing datasets was performed by the BRC Genomics Core at KCL using Cell Ranger-4.0.0. Once feature-barcode matrices were obtained, analysis was performed in RStudio with the Seurat package, v3 and 4 ^36,37^, following author instructions. The dataset was subset to exclude cells with <500 or >5000 genes or with >20% mitochondrial genes. After normalisation, the 2000 most variable features were identified, the dataset was scaled and PCA was estimated based on the previously identified variable features. A UMAP was generated using the top 10 PCs and a resolution of 0.4. Cluster identities were assigned based on markers from the literature, gene counts from clusters showing medulla-specific markers were extracted and the matrix re-analysed with the same parameters. Cell cycle analysis was performed using the CellCycleScoring function in Seurat, following author specifications. Pseudotime analysis was performed using Monocle ^33^, following author instructions.

### 10x Single-cell RNA sequencing and computational analysis - postnatal SOX2-EGFP^+^ cells

30 adrenals from 15 mixed sex P15 S*ox2^eGFP/+^* mice were dissected on ice, dissociated, and GFP+ cells were isolated via FACS, centrifuged at 300g for 5 minutes and resuspended in a solution of HBSS 2.5% FBS. 2,000 viable cells were used for the experiment. Droplet-based single-cell RNA sequencing was performed as described. Cells with <1000 or >5000 genes or with >20% mitochondrial genes were excluded. After normalisation, the 2000 most variable features were identified, the dataset was scaled and PCA was estimated based on the previously identified variable features. A UMAP was generated using the top 10 PCs and a resolution of 0.4. To further select only *Sox2* expressing cells, the WhichCell function was used to select only cells with *Sox2* normalised expression >1. Once extracting the raw counts from these cells, the dataset was reanalysed with the same parameters.

### Differential expression analysis of SCPs vs *Sox2* expressing postnatal cells

SCPs were isolated from a 13.5dpc dataset^15^ using the parameters described in the paper. This dataset was combined with the *Sox2* expressing cells dataset using Seurat integration PCs 1:30 and 2000 variable features. Differential expression analysis between SCPs and postnatal *Sox2* expressing cells was performed following Seurat guidelines. https://www.zotero.org/google-docs/?XvRWI5ClusterProfiler^34^ was used to obtain significantly differentially expressed gene ontologies. STRING https://www.zotero.org/google-docs/?tmJnpdanalysis^35^ of the 50 top differentially expressed genes in the postnatal dataset was used to reveal connections.

### Correlation analysis of Sox2 regulon

Data published https://www.zotero.org/google-docs/?nqcKLJby^29^ were utilised. This dataset included the processed transcriptomic data as well as the SCENIC-related dataset. The Spearman correlation between the activity score of the Sox2(+) regulon and the log10 expression of gene transcripts in all processed cells was calculated. Expression values of top correlated and anti-correlated genes were also represented by fitting these with Generalized-Additive model (GAM), using the pseudotime assignments of the cells of the trajectory from late NCC/SCP to ChC. Pseudotime and trajectory representation and analysis were carried out using scFates package.

### Tissue processing

For paraffin-embedding, adrenal glands were dissected, surrounding fat was removed and samples were fixed in 10% neutral buffered formalin (NBF) (Sigma, HT501128) overnight at room temperature. Grafts collected from the chorioallantoic membranes (CAM) were dissected and fixed following the same protocol. The next day, tissue was washed and dehydrated through graded ethanol series and paraffin-embedded. Samples were sectioned at 5μm thickness. For cryo-embedding, adrenal glands were dissected, surrounding fat removed and samples fixed in 4% PFA at 4°C for 4 hours. Adrenals were washed and cryoprotected in 30% Sucrose overnight at 4°C. Adrenals were embedded in Optical Cutting Temperature (OCT) compound (VWR, 361603E) and flash-frozen. Samples were cryo- sectioned at 8-12μm thickness.

### Immunofluorescence and immunohistochemistry on paraffin sections

Paraffin sections were deparaffinised and rehydrated with ethanol series. Antigen retrieval was performed in a Decloaking Chamber NXGEN (Menarini Diagnostics, DC2012-220V) at 110°C for 3 minutes using Declere, pH 6.0 (Cell Marque, 921P-04) for immunohistochemistry or Dako Target Retrieval Solution, pH 9.0 (Agilent, S236784-2) for immunofluorescence.

For immunohistochemistry, ImmPRESS Excel Amplified HRP Polymer Staining Kit Anti-Rabbit IgG (Vector Laboratories, MP-7602-50) was used following the manufacturer’s instructions. Primary antibodies were used at the concentrations listed in STAR Methods Resource Table. Nuclei were stained with Vector Hematoxylin QS (Vector Laboratories, H-3404-100) and slides were mounted in VectaMount Permanent Mounting Medium (Vector Laboratories, H-5000-60).

For immunofluorescence, sections were blocked for 1 hour at room temperature in Blocking Buffer (0.15% glycine, 2mg/ml BSA, 0.1% Triton X-100 in PBS) with 10% sheep serum. Primary antibodies were diluted in Blocking Buffer with 1% sheep serum at the concentrations described in STAR Methods

Resource Table and incubated overnight at 4°C. After washing 3 times with PBST (PBS + 0.1% Triton X- 100), samples were incubated for 1 hour at room temperature in secondary fluorophore-conjugated antibodies (dilution 1:500, listed in STAR Methods Resource Table) and Hoechst (Life Technologies, H3570) (dilution 1:10,000) in blocking buffer with 1% serum. Tyrosine Hydroxylase and PNMT antibodies were amplified with biotin-streptavidin by incubating at room temperature for 1 hour with anti-mouse biotinylated secondary antibody (dilution 1:300, listed in STAR Methods Resource Table) and Hoechst (Life Technologies, H3570) (dilution 1:10,000), washed 3 times with PBST and incubated at room temperature for 1 hour with fluorescent-labelled streptavidin (dilution 1:500, listed in STAR Methods Resource Table). After washing in PBST, slides were mounted with Vectashield Antifade Mounting Medium (Vector Laboratories, H-1000-10).

### Immunofluorescence on cryosections

Sections were blocked for 1 hour at room temperature in Blocking Buffer (1% BSA, 0.1% Triton X-100, 5% goat serum). Primary antibodies were diluted in Blocking Buffer at the concentrations reported in STAR Methods Resource Table and incubated overnight at 4°C. After washing 3 times with PBS, secondary fluorophore-conjugated antibodies (dilution 1:500, listed in STAR Methods Resource Table) and Hoechst (Life Technologies, H3570) (dilution 1:10,000) were diluted in in Blocking Buffer and incubated for 1 hour at room temperature. After washing 3 times with PBS, slides were mounted with Vectashield Antifade Mounting Medium (Vector Laboratories, H-1000-10).

### RNAscope mRNA *in situ* hybridisation

RNAscope was performed on paraffin-embedded sections with the RNAscope 2.5 HD Duplex Kit (ACD Bio, 322430) following the manufacturer’s protocol, with optimised retrieval time of 12 minutes and protease time of 30 minutes. Probes used are listed in STAR Methods Resource Table. Sections were counterstained with Hematoxylin QS (Vector Laboratories, H-3404-100) and slides were mounted in VectaMount Permanent Mounting Medium (Vector Laboratories, H-5000-60).

### Primary cell culture

Adrenal glands were dissected, and the medulla isolated manually. Medullae were placed in an enzymatic digestion mix containing 50μg/ml DNAse I (Sigma, D5025), 10mg/ml Collagenase II (Worthington, LS004177), 2.5μg/ml Fungizone (Gibco, 15290026), 0.1X Trypsin-EDTA (Sigma, 59418C) in 1X Hank’s Balanced Salt Solution (HBSS) (Gibco, 14025050). Medullae in enzymatic digestion mix were incubated at 37°C for 10 minutes, triturated by pipetting up and down and incubated for 5 minutes at 37°C, followed by trituration to obtain a single-cell suspension. Enzymes were inactivated by addition of 10 times volume of serum-containing Base Media: DMEM/F-12 (Gibco, 31330-038) + 5% FBS (Merk, F0804) + 50U/ml Penicillin-Streptomycin (Gibco, 15070063). The cell suspension was centrifuged at 300g for 5 minutes at room temperature, washed twice in PBS before being resuspended in Complete Media: Base Media + 20ng/ml bFGF (R&D Systems, 234-FSE) + 50μg/ml cholera toxin (Sigma, C8052). Two days after isolation, an equal volume of fresh media was added to each plate. Media was fully changed every 2-3 days. For immunostaining, cells were plated on glass coverslips coated with 0.1% gelatine diluted in PBS.

### Colony Forming Assay

For colony forming assays, adrenals from *Sox2^eGFP/+^* mice were dissected and tissue was dissociated as described for primary cell culture. GFP^+^ and GFP^-^ cells were separated by flow sorting. After sorting, GFP^+^ and GFP^-^ cells were plated at clonal density of 500 cells/well in a 12-well plate. Two days after isolation, an equal volume of medium to the one present in the plate was added. After that, media were changed every 2-3 days.

After 14 days of culture, cells were washed 3 times in PBS and fixed with 10% NBF for 10 minutes at room temperature. After washing 3 times with PBS, cells were stained for 10 minutes with Crystal Violet Solution: 0.5% Crystal Violet powder (Sigma, C0775), 20% methanol in distilled water. Excess crystal violet was washed with running tap water and plates dried before colony observation and imaging.

### Chorioallantoic Membrane (CAM) assays

Fertilised Shaver Brown eggs were purchased from Medeggs Ltd and placed in an egg incubator set at 37.8°C/60% humidity. This is considered developmental day 0. On day 4, the CAM was exposed using curved spring scissors and the window sealed with clear tape to prevent contamination and placed back in the incubator. On day 10 of incubation, a silicone ring of 6mm diameter was placed onto the CAM of each egg. 8 x 10^5^ isolated *Sox2^eGFP^* cells were seeded within the silicone ring. The window was sealed again and the eggs placed in the incubator until graft collection at day 18.

### Immunofluorescence on cells

For immunofluorescence on cells, coverslips were washed twice in PBS and fixed with 4% PFA on ice for 10 minutes. After washing with PBST, cells were blocked for 1 hour at room temperature in Blocking Buffer (0.15% glycine, 2mg/ml BSA, 0.1% Triton X-100 in PBS) with 10% sheep serum. Primary antibodies were incubated overnight at 4°C in Blocking Buffer with 1% sheep serum at the concentrations shown in the STAR Methods Resource Table. After washing with PBST, sections were incubated for 1 hour at room temperature in secondary fluorophore-conjugated antibodies, diluted 1:500 (listed in STAR Methods Resource Table) in Blocking Buffer with 1% serum. After washing with PBST, coverslips were mounted with Vectashield HardSet Antifade Mounting Medium with DAPI (Vector Laboratories, H-1500-10)

### Imaging

Images of immunofluorescence staining were taken with a Leica TCS SP5 or a Zeiss LSM980 confocal microscope, using an HCX Plan-Apochromat CS 20x/0.7 dry objective and an HCX Plan-Apochromat CS 63x/1.3 Glycine objective (both Leica Microsystems), or Zeiss Plan-Apochromat 20x/0.8 dry objective, a Zeiss C-Apochromat 40x/1.2 Water objective and a Zeiss Plan-Apochromat 63x/1.40 Oil objective.

Stacks of 0.5μm/0.7μm were taken through the entire section thickness. Immunohistochemistry and RNAscope stained sections were scanned with a Nanozoomer-XR Digital slide scanner (Hamamatsu), close-up images were taken with an Olympus BX34F Brightfield microscope using a 40X objective. Cell culture images were taken with an Olympus Phase Contrast microscope using a 4X or 20X objective. Images were processed with Fiji (Schindelin et al., 2012) and with Nanozoomer Digital Pathology View. Figures were created in Adobe Illustrator.

### Quantifications and statistics

Cell counting was performed manually with Fiji’s “Cell Counter” plugin. For mouse samples, a minimum of three sections per sample were counted. For human samples, 5 representative fields were selected at 20X magnification for each sample and counted. Statistical analysis and graphs (except for single-cell RNA sequencing analysis) were produced using GraphPad Prism.

### Code and data availability

Code is available at: https://github.com/Andoniadou-Lab/adrenal_stemcell Datasets are available at the Gene Expression Omnibus (GEO) with accession number GSE237125.

---

**Figure S1.**
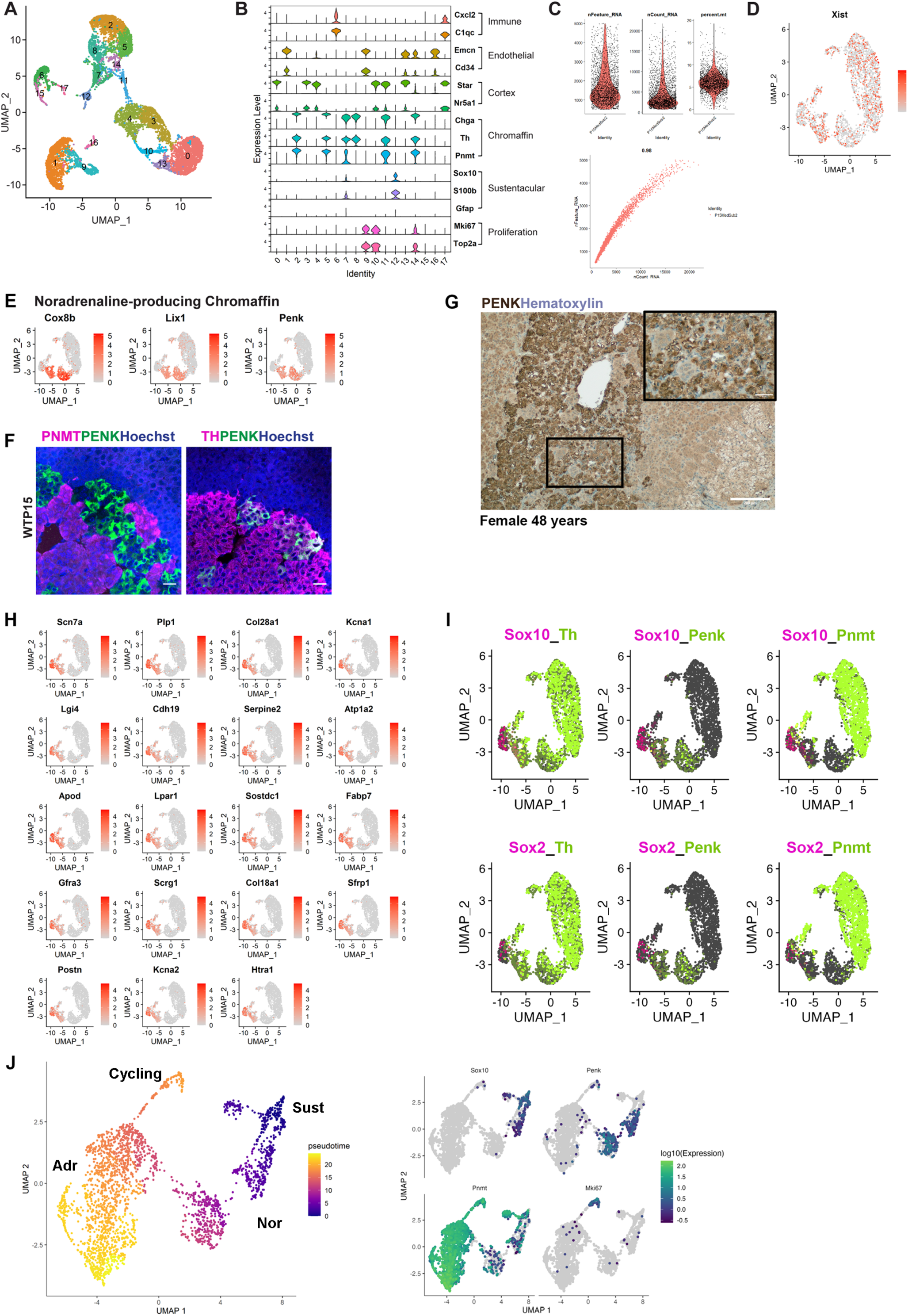
Single-cell RNA sequencing of the mouse adrenal medulla. A) UMAP of the entire postnatal medulla dataset obtained (9961 cells); B) Violin plots indicating expression markers chosen for downstream subsetting. C) QC for final dataset used. D) Featureplot for expression of *Xist*, indicating female cells in the dataset. E) Featureplots showing newly identified markers *Cox8b*, *Lix1* and *Penk*, specific to the noradrenaline-producing chromaffin cell cluster. F) Immunofluorescence with antibodies against PNMT or TH (magenta) and against PENK (green)on wild type P15 adrenals. Nuclei counterstained with Hoechst, scale bar 20μm. G) Immunohistochemistry on a human adrenal medulla (Female, 48 years of age) using antibodies against PENK (brown) confirming expression. Nuclei counterstained with Hematoxylin, Scale bar 200μm, inset 50μm. H) Featureplots showing newly identified markers of mouse sustentacular cells. I) Featureplots showing gene expression of *Sox2* or *Sox10* with differentiated cell markers *Th*, *Penk*, *Pnmt*. J) Monocle pseudotime UMAP, marker gene expression plotted on Monocle UMAP. Sust- sustentacular cells; Nor- Noradrenaline chromaffin lineage; Adr- Adrenaline chromaffin lineage; Cycling- cycling adrenaline chromaffin cells.

**Figure S2.**
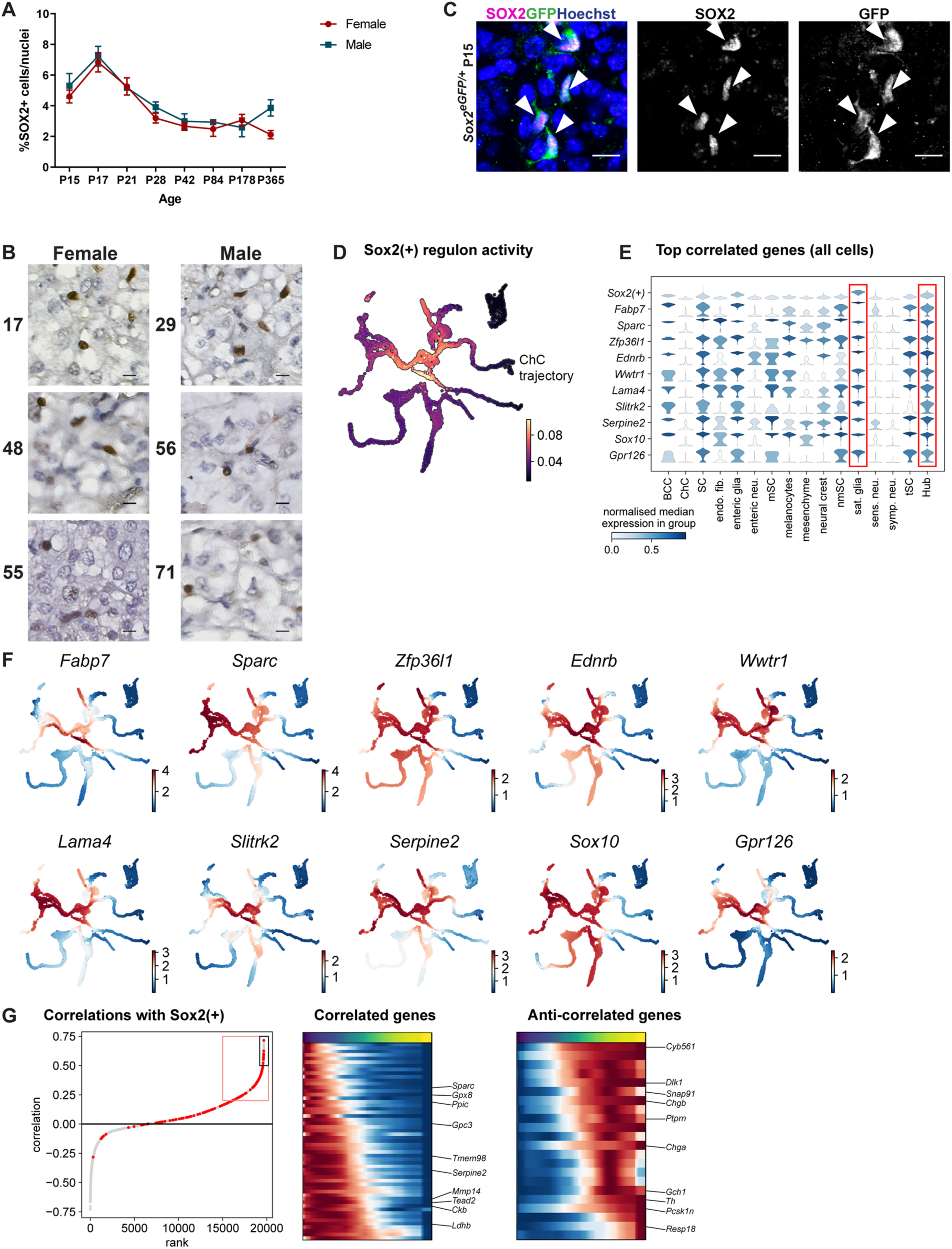
SOX2^+^ cells are present in the mouse adrenal medulla and are derived from Schwann Cell Precursors. A) Quantification of SOX2^+^ cells over the total nuclei of adrenal medulla, split by sex. n = 3 animals per sex, per timepoint. Mean and SEM plotted. B) Immunohistochemistry with antibodies against SOX2 (brown) on normal human adrenals in females (at 17, 48 and 55 years of age) and males (at 29, 56, and 71 years of age). Nuclei counterstained with Hematoxylin, scale bar 20μm. C) Immunofluorescence on *Sox2^eGFP/+^* adrenal medulla at P15, using antibodies against GFP (green) and SOX2 (magenta) showing complete co-localisation (arrowheads). Nuclei counterstained with Hoechst, scale bars 10μm. D) Activity of the *Sox2* regulon in data from ^29^, showing this is active in the early part of the chromaffin cell trajectory (labelled). E) Top correlated genes to *Sox2*, irrespective of trajectory, which include several markers of the postnatal *Sox2*-expressing population. The highest expression is observed in satellite glia and Hub cells (red boxes). F) Featureplots of *Sox2*-correlated genes expression. G) Ranked correlation analysis identifying genes highest correlated with *Sox2* specifically in the chromaffin cell trajectory, and anti-correlated with *Sox2*, which includes chromaffin cell markers (*Th*, *Chga*, *Chgb*).

**Figure S3.**
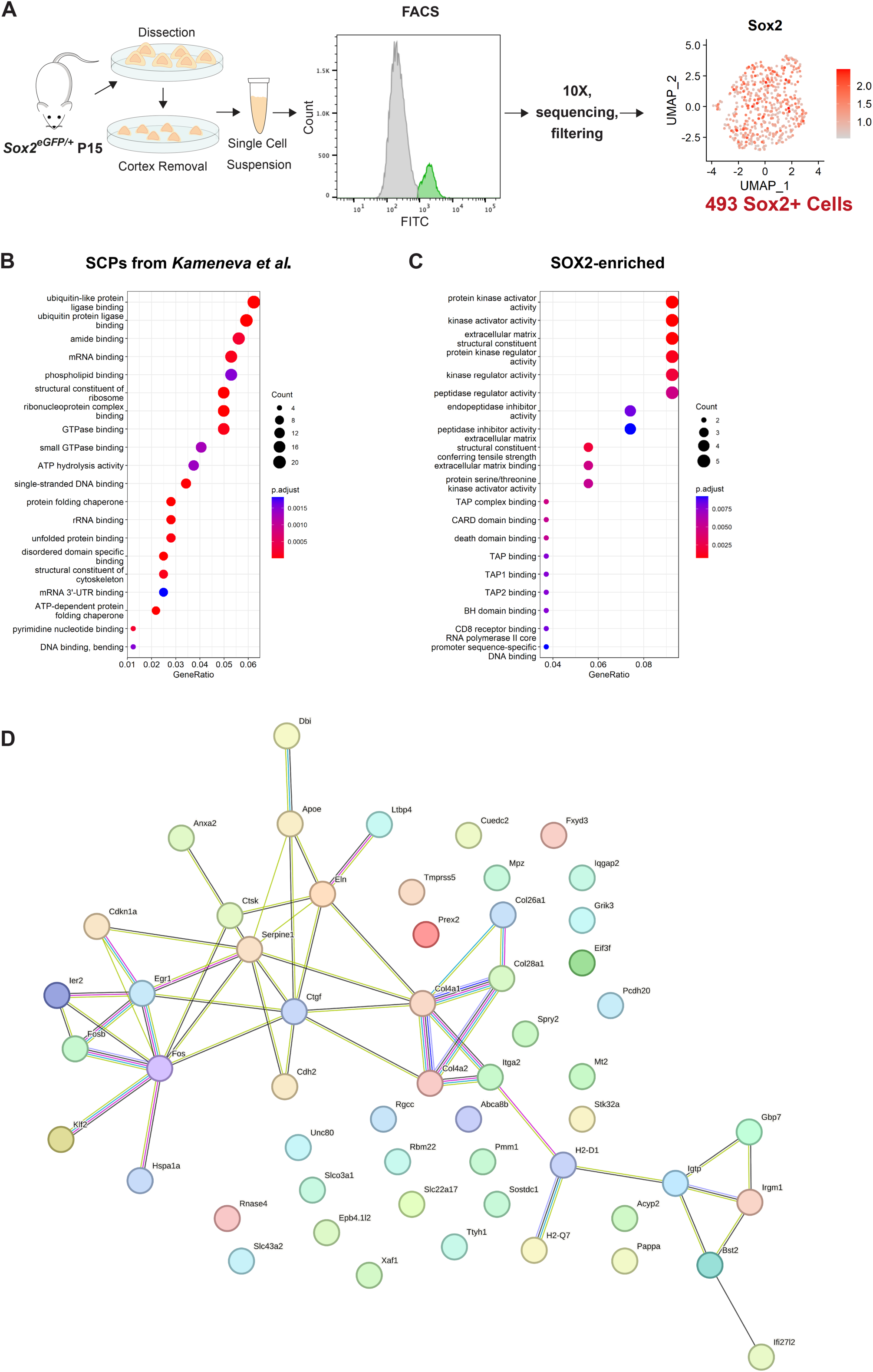
Adrenomedullary SOX2^+^ stem cells are a distinct population from Schwann Cell Precursors. A) Experimental design. Featureplot showing *Sox2* expression in the selected dataset. B) Gene ontology of differentially expressed genes by embryonic SCPs compared to postnatal *Sox2*-expressing cells. C) Gene ontology of differentially expressed genes by postnatal *Sox2*-expressing cells compared to embryonic SCPs. D) STRING analysis of the top 50 most differentially expressed genes by postnatal *Sox2*-expressing cells compared to embryonic SCPs.

**Figure S4.**
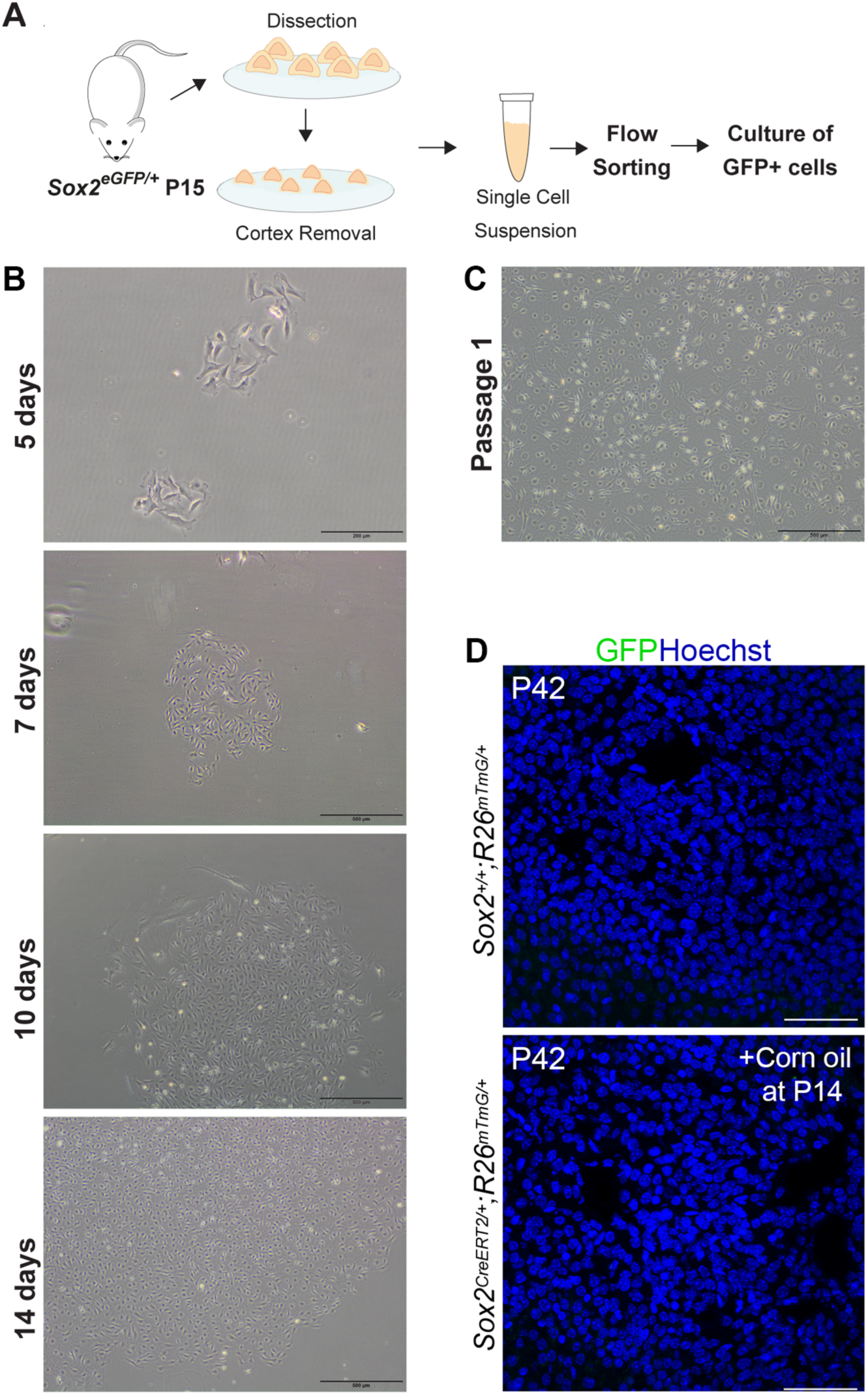
Adrenomedullary SOX2^+^ cells have stem cell properties. A) Experimental design. B) Brightfield images of cultured GFP+ cells: 5, 7, 10, 14 days after isolation. Scale bars 200μm (5 days) and 500μm (7, 10, 14 days). C) Brightfield image of cultured GFP+ cells after 1 passage, scale bar 500μm. D) Immunofluorescence using antibodies against GFP (green) on sections from a *Sox2^+/+^;R26^mTmG/+^*adrenal at P42 (top panel) or *Sox2^CreERT^*^2^*^/+^;R26^mTmG/+^* adrenal from a mouse injected with corn oil P14 and collected after 28 days (P42, bottom panel). Note the absence of GFP-labelled cells in these controls. Nuclei counterstained with Hoechst (blue), scale bar 50μm.

**Figure S5.**
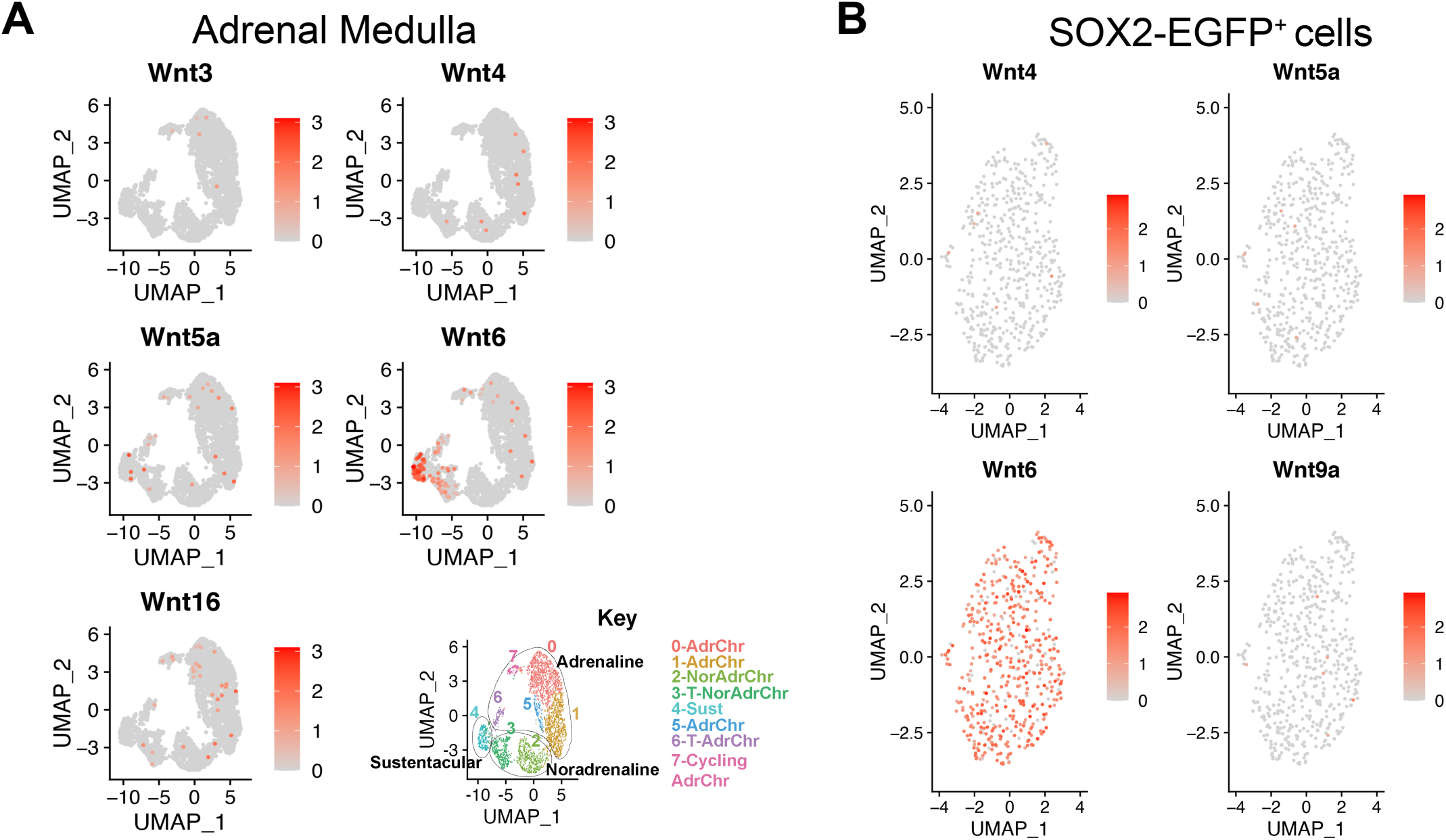
SOX2+ adrenomedullary stem cells promote proliferation of chromaffin cells through secretion of paracrine WNT signals. A) Featureplots for *Wnt3*, *Wnt4*, *Wnt5a*, *Wnt6*, and *Wnt16* in the mouse adrenal medulla dataset. B) Featureplots for *Wnt4*, *Wnt5a*, *Wnt6*, and *Wnt9a* in the isolated SOX2-EGFP^+^ cell dataset.

## References

1. Kvetnansky, R., Sabban, E.L., and Palkovits, M. (2009). Catecholaminergic systems in stress: structural and molecular genetic approaches. Physiol. Rev. 89, 535–606. 10.1152/physrev.00042.2006.

2. Kvetnansky, R., Lu, X., and Ziegler, M.G. (2013). Stress-triggered changes in peripheral catecholaminergic systems. Adv. Pharmacol. San Diego Calif 68, 359–397. 10.1016/B978-0-12-411512-5.00017-8.

3. Bechmann, N., Berger, I., Bornstein, S.R., and Steenblock, C. (2021). Adrenal medulla development and medullary-cortical interactions. Mol. Cell. Endocrinol. 528, 111258. 10.1016/j.mce.2021.111258.

4. Kim, M.S., Ryabets-Lienhard, A., Bali, B., Lane, C.J., Park, A.H., Hall, S., and Geffner, M.E. (2014). Decreased adrenomedullary function in infants with classical congenital adrenal hyperplasia. J. Clin. Endocrinol. Metab. 99, E1597–1601. 10.1210/jc.2014-1274.

5. Lenders, J.W.M., Eisenhofer, G., Mannelli, M., and Pacak, K. (2005). Phaeochromocytoma. Lancet Lond. Engl. 366, 665–675. 10.1016/S0140-6736(05)67139-5.

6. Bedoya-Reina, O.C., Li, W., Arceo, M., Plescher, M., Bullova, P., Pui, H., Kaucka, M., Kharchenko, P., Martinsson, T., Holmberg, J., et al. (2021). Single-nuclei transcriptomes from human adrenal gland reveal distinct cellular identities of low and high-risk neuroblastoma tumors. Nat. Commun. 2021 121 12, 1–15. 10.1038/s41467-021-24870-7.

7. Hanemaaijer, E.S., Margaritis, T., Sanders, K., Bos, F.L., Candelli, T., Al-Saati, H., van Noesel, M.M., Meyer-Wentrup, F.A.G., van de Wetering, M., Holstege, F.C.P., et al. (2021). Single-cell atlas of developing murine adrenal gland reveals relation of Schwann cell precursor signature to neuroblastoma phenotype. Proc. Natl. Acad. Sci. U. S. A. 118. 10.1073/pnas.2022350118.

8. Lopez, J.P., Brivio, E., Santambrogio, A., Donno, C.D., Kos, A., Peters, M., Rost, N., Czamara, D., Brückl, T.M., Roeh, S., et al. (2021). Single-cell molecular profiling of all three components of the HPA axis reveals adrenal ABCB1 as a regulator of stress adaptation. Sci. Adv. 7, eabe4497. 10.1126/SCIADV.ABE4497.

9. Chung, K.F., Sicard, F., Vukicevic, V., Hermann, A., Storch, A., Huttner, W.B., Bornstein, S.R., and Ehrhart-Bornstein, M. (2009). Isolation of neural crest derived chromaffin progenitors from adult adrenal medulla. Stem Cells 27, 2602–2613. 10.1002/stem.180.

10. Rubin De Celis, M.F., Garcia-Martin, R., Wittig, D., Valencia, G.D., Enikolopov, G., Funk, R.H., Chavakis, T., Bornstein, S.R., Androutsellis-Theotokis, A., and Ehrhart-Bornstein, M. (2015). Multipotent glia-like stem cells mediate stress adaptation. Stem Cells 33, 2037–2051. 10.1002/stem.2002.

11. Santana, M.M., Chung, K.-F., Vukicevic, V., Rosmaninho-Salgado, J., Kanczkowski, W., Cortez, V., Hackmann, K., Bastos, C.A., Mota, A., Schrock, E., et al. (2012). Isolation, Characterization, and Differentiation of Progenitor Cells from Human Adult Adrenal Medulla. STEM CELLS Transl. Med. 1, 783–791. 10.5966/sctm.2012-0022.

12. Tischler, A.S., Ruzicka, L.A., Donahue, S.R., and DeLellis, R.A. (1989). Chromaffin cell proliferation in the adult rat adrenal medulla. Int. J. Dev. Neurosci. 7, 439–448. 10.1016/0736-5748(89)90004-X.

13. Verhofstad, A.A. (1993). Kinetics of adrenal medullary cells. J. Anat. 183 (Pt 2), 315–326.

14. Furlan, A., Dyachuk, V., Kastriti, M.E., Calvo-Enrique, L., Abdo, H., Hadjab, S., Chontorotzea, T., Akkuratova, N., Usoskin, D., Kamenev, D., et al. (2017). Multipotent peripheral glial cells generate neuroendocrine cells of the adrenal medulla. Science 357, eaal3753. 10.1126/science.aal3753.

15. Kameneva, P., Artemov, A.V., Kastriti, M.E., Faure, L., Olsen, T.K., Otte, J., Erickson, A., Semsch, B., Andersson, E.R., Ratz, M., et al. (2021). Single-cell transcriptomics of human embryos identifies multiple sympathoblast lineages with potential implications for neuroblastoma origin. Nat. Genet. 53, 694–706. 10.1038/s41588-021-00818-x.

16. Lumb, R., Tata, M., Xu, X., Joyce, A., Marchant, C., Harvey, N., Ruhrberg, C., and Schwarz, Q. (2018). Neuropilins guide preganglionic sympathetic axons and chromaffin cell precursors to establish the adrenal medulla. Development 145, dev.162552. 10.1242/dev.162552.

17. Bielohuby, M., Herbach, N., Maser-gluth, C., Beuschlein, F., Wolf, E., and Hoeflich, A. (2007). Growth analysis of the mouse adrenal gland from weaning to adulthood: time- and gender-dependent alterations of cell size and number in the cortical compartment. Am J Physiol Endocrinol Metab 293 *293*, E139–E146. 10.1152/ajpendo.00705.2006.

18. Viveros, O.H., Diliberto, E.J., Hazum, E., and Chang, K.J. (1979). Opiate-like materials in the adrenal medulla: Evidence for storage and secretion with catecholamines. Mol. Pharmacol. 16, 1101–1108.

19. Takahashi, M., and Osumi, N. (2005). Identification of a novel type II classical cadherin: Rat cadherin19 is expressed in the cranial ganglia and Schwann cell precursors during development. Dev. Dyn. 232, 200–208. 10.1002/dvdy.20209.

20. Andoniadou, C.L., Matsushima, D., Neda, S., Gharavy, M., Signore, M., Mackintosh, A.I., Schaeffer, M., Gaston-Massuet, C., Mollard, P., Jacques, T.S., et al. (2013). Sox2+ stem/progenitor cells in the adult mouse pituitary support organ homeostasis and have tumor-inducing potential. Cell Stem Cell 13, 433–445. 10.1016/j.stem.2013.07.004.

21. Ellis, P., Fagan, B.M., Magness, S.T., Hutton, S., Taranova, O., Hayashi, S., McMahon, A., Rao, M., and Pevny, L. (2004). SOX2, a persistent marker for multipotential neural stem cells derived from embryonic stem cells, the embryo or the adult. Dev. Neurosci. 26, 148–165. 10.1159/000082134.

22. Emmerson, E., May, A.J., Nathan, S., Cruz-Pacheco, N., Lizama, C.O., Maliskova, L., Zovein, A.C., Shen, Y., Muench, M.O., and Knox, S.M. (2017). SOX2 regulates acinar cell development in the salivary gland. eLife 6. 10.7554/eLife.26620.

23. Que, J., Luo, X., Schwartz, R.J., and Hogan, B.L.M. (2009). Multiple roles for Sox2 in the developing and adult mouse trachea. Development 136, 1899 LP – 1907. 10.1242/dev.034629.

24. Auerbach, R., Kubai, L., Knighton, D., and Folkman, J. (1974). A simple procedure for the long-term cultivation of chicken embryos. Dev. Biol. 41, 391–394. 10.1016/0012-1606(74)90316-9.

25. Russell JP, Lim X, Santambrogio A, Yianni V, Kemkem Y, Wang B, Fish M, Haston S, Grabek A, Hallang S, Lodge EJ, Patist AL, Schedl A, Mollard P, Nusse R, Andoniadou CL. (2021). Pituitary stem cells produce paracrine WNT signals to control the expansion of their descendant progenitor cells. elife. 5. 10.7554/eLife.59142.

26. François, A., Low, S.A., Sypek, E.I., Christensen, A.J., Sotoudeh, C., Beier, K.T., Ramakrishnan, C., Ritola, K.D., Sharif-Naeini, R., Deisseroth, K., et al. (2017). A Brainstem-Spinal Cord Inhibitory Circuit for Mechanical Pain Modulation by GABA and Enkephalins. Neuron 93, 822–839.e6. 10.1016/j.neuron.2017.01.008.

27. Kastriti, M.E., Kameneva, P., and Adameyko, I. (2020). Stem cells, evolutionary aspects and pathology of the adrenal medulla: A new developmental paradigm. Mol. Cell. Endocrinol. 518, 110998. 10.1016/J.MCE.2020.110998. 11. 10.3389/fendo.2020.00079.

28. Scriba, L.D., Bornstein, S.R., Santambrogio, A., Mueller, G., Huebner, A., Hauer, J., Schedl, A., Wielockx, B., Eisenhofer, G., Andoniadou, C.L., et al. (2020). Cancer Stem Cells in Pheochromocytoma and Paraganglioma. Front. Endocrinol. 11. 10.3389/fendo.2020.00079.

29. Kastriti, M.E., Faure, L., Von Ahsen, D., Bouderlique, T.G., Boström, J., Solovieva, T., Jackson, C., Bronner, M., Meijer, D., Hadjab, S., et al. (2022). Schwann cell precursors represent a neural crest-like state with biased multipotency. EMBO J. 41, e108780. 10.15252/embj.2021108780.

30. Danielian, P.S., Muccino, D., Rowitch, D.H., Michael, S.K., and McMahon, A.P. (1998). Modification of gene activity in mouse embryos in utero by a tamoxifen-inducible form of Cre recombinase. Curr. Biol. CB.

31. Laranjeira, C., Sandgren, K., Kessaris, N., Richardson, W., Potocnik, A., Vanden Berghe, P., and Pachnis, V. (2011). Glial cells in the mouse enteric nervous system can undergo neurogenesis in response to injury. J. Clin. Invest. 121, 3412–3424. 10.1172/JCI58200.

32. Muzumdar, M.D., Tasic, B., Miyamichi, K., Li, L., and Luo, L. (2007). A Global Double-Fluorescent Cre Reporter Mouse. genesis 45, 593–605. 10.1002/dvg.

33. Trapnell, C., Cacchiarelli, D., Grimsby, J., Pokharel, P., Li, S., Morse, M., Lennon, N.J., Livak, K.J., Mikkelsen, T.S., and Rinn, J.L. (2014). The dynamics and regulators of cell fate decisions are revealed by pseudotemporal ordering of single cells. Nat. Biotechnol. 32, 381–386. 10.1038/nbt.2859.

34. Wu, T., Hu, E., Xu, S., Chen, M., Guo, P., Dai, Z., Feng, T., Zhou, L., Tang, W., Zhan, L., et al. (2021). clusterProfiler 4.0: A universal enrichment tool for interpreting omics data. The Innovation 2. 10.1016/j.xinn.2021.100141.

35. Szklarczyk, D., Franceschini, A., Wyder, S., Forslund, K., Heller, D., Huerta-Cepas, J., Simonovic, M., Roth, A., Santos, A., Tsafou, K.P., et al. (2015). STRING v10: protein-protein interaction networks, integrated over the tree of life. Nucleic Acids Res. 43, D447–452. 10.1093/nar/gku1003.

36. Hao, Y., Hao, S., Andersen-Nissen, E., Mauck, W.M., Zheng, S., Butler, A., Lee, M.J., Wilk, A.J., Darby, C., Zager, M., et al. (2021). Integrated analysis of multimodal single-cell data. Cell 184, 3573–3587.e29. 10.1016/J.CELL.2021.04.048.

37. Stuart, T., Butler, A., Hoffman, P., Hafemeister, C., Papalexi, E., Mauck, W.M., Hao, Y., Stoeckius, M., Smibert, P., and Satija, R. (2019). Comprehensive Integration of Single-Cell Data. Cell 177, 1888–1902.e21. 10.1016/J.CELL.2019.05.031.

